# Targeting matrix metalloproteinase-14 disrupts DNA repair and reduces viability in adrenocortical carcinoma

**DOI:** 10.64898/2026.01.06.697992

**Authors:** Changxian Shen, Liudmila V. Popova, Daniel M. Chopyk, Lindsey Hartshorn, Zhongguang Li, Yaoling Shu, Caleb Araujo, Shivam Priya, Varsha Thakur, John E. Phay, Barbra S. Miller, Barbara Bedogni, Haichang Li, Priya H. Dedhia

**Affiliations:** Division of Surgical Oncology, The Ohio State University and Arthur G. James Comprehensive Cancer Center; Columbus, Ohio, USA; Department of Veterinary Biosciences, College of Veterinary Medicine, The Ohio State University, Columbus, Ohio, USA; Department of Cancer Biology and Genetics, College of Medicine, The Ohio State University; Columbus, Ohio, USA; Dr. Phillip Frost Department of Dermatology and Cutaneous Surgery, University of Miami Miller School of Medicine, Sylvester Comprehensive Cancer Center, Miami, Florida, USA; Division of Surgical Oncology, University of Arizona College of Medicine, Tucson, AZ; Translational Therapeutics Program, The Ohio State University and Arthur G. James Comprehensive Cancer Center; Columbus, Ohio, USA; Center for Cancer Engineering, The Ohio State University; Columbus, Ohio, USA

**Keywords:** Adrenocortical carcinoma, matrix metalloproteinase-14, hemopexin domain, DNA replication stress, DNA repair, patient derived tumor organoid

## Abstract

**Background:** Adrenocortical carcinoma (ACC) is a rare and aggressive endocrine cancer with limited treatment options and poor prognosis. Identifying novel therapeutic targets requires understanding the molecular drivers of ACC progression and establishing translational models for preclinical validation.

**Results:** Matrix metalloproteinase-14 (MMP-14) is the most highly expressed MMP in ACC, and high MMP-14 expression is associated with worse overall and disease-free survival. We demonstrate that MMP-14 is essential for ACC cell survival and serves an unexpected role in maintaining genome stability. Genetic silencing or pharmacologic inhibition of MMP-14 significantly reduced viability in both NCI-H295R cells and patient-derived tumor organoids (PTOs). MMP-14 knockdown induced CHK1 activation and S-phase checkpoint arrest. Mechanistically, MMP-14 translocates to the nucleus and binds to chromatin following DNA damage induced by ionizing radiation or cisplatin. Loss of MMP-14 resulted in accumulation of DNA double-strand breaks, as evidenced by increased γH2AX foci, and impaired non-homologous end joining (NHEJ)-mediated repair.

**Conclusions:** These findings reveal a novel nuclear function for MMP-14 in DNA repair and identify MMP-14 as a promising therapeutic target in ACC. Targeting MMP-14 may sensitize ACC tumors to DNA-damaging chemotherapy by impairing the repair of therapy-induced lesions.

## BACKGROUND

Adrenocortical carcinoma (ACC) is a rare yet highly aggressive malignancy of the adrenal cortex. The majority of patients present with advanced stage disease, for which effective therapeutic options remain limited[1]. Poor survival outcomes underscore the urgent need for novel therapeutic targets. Current first-line treatment-mitotane combined with etoposide, doxorubicin, and cisplatin-extends progression-free survival by only 5 months with response rates of approximately 23%[2]. Targeted therapies-including IGF-1R and multikinase inhibitors-and immunotherapy have shown limited efficacy[3–6]. These therapeutic failures highlight the critical need to identify novel molecular targets in ACC.

Matrix metalloproteinases (MMPs) remodel the extracellular matrix (ECM) through proteolytic degradation of collagen and other ECM components, and can contribute to cancer progression through cell migration, invasion, and angiogenesis[7]. MMP-14, also known as membrane type matrix metalloproteinase 1 (MT-MMP1), is distinct from secreted MMPs in that it is anchored to the plasma membrane, enabling spatially restricted proteolysis at the cell surface[3]. This membrane localization allows MMP-14 to proteolytically activate other pro-MMPs and pro-proteins[7, 8], thereby altering the tumor microenvironment through release of embedded growth factors, cytokines, and their receptors into soluble, active signals that influence nearby cells[9–11].

MMP-14 promotes cell migration and invasion via multiple mechanisms. Specifically, MMP-14 localizes to invadopodia-actin-rich membrane protrusions that mediate focused ECM degradation-where the protein directs matrix breakdown to enable cell migration[7]. Beyond ECM remodeling, MMP-14 can induce an epithelial-to-mesenchymal transition (EMT)-like phenotype and facilitate cell motility through cleavage of cell-cell adhesion molecules such as E-cadherin and indirect upregulation of mesenchymal markers (e.g., vimentin). Additionally,

MMP-14 can be packaged into exosomes and transferred to neighboring cells, promoting migration and conferring chemoresistance in recipient cells[12]. MMP-14 also promotes angiogenesis through two mechanisms: 1) degradation of the vascular basement membrane to guide endothelial cell migration and sprouting, and 2) activation of pro-angiogenic factors such as (VEGF) and Heparin-Binding EGF-like Growth Factor (HB-EGF) through proteolytic cleavage[7, 13].

The diverse functions of MMP-14 are mediated by its distinct structural domains. MMP-14 contains a catalytic domain, a hemopexin domain, a transmembrane domain, and a cytoplasmic tail[14]. Beyond proteolysis by the catalytic domain, the MMP-14 cytoplasmic tail engages in non-proteolytic signaling, which can activate intracellular signaling pathways in tumor cells such as ERK/MAPK, PI3K/AKT, Src, and RhoA/Rac1[9, 15–18]. While canonical ECM-related functions are well established, emerging evidence suggests MMP-14 may have non-canonical roles in DNA replication[19]. However, the precise underlying molecular mechanisms and the biological context of these functions remain largely unknown[20].

Consistent with pro-tumorigenic functions, MMP-14 overexpression occurs frequently across multiple cancer types and correlates with tumor progression and poor prognosis[21–24]. In ACC specifically, MMP-14 expression is elevated, and patients with higher MMP-14 levels demonstrate worse survival outcomes[24]. Despite this clinical association, the therapeutic potential of MMP-14 inhibition in ACC has not been evaluated.

Given the elevated expression of MMP-14 in ACC and its poorly understood non-canonical functions, we investigated the therapeutic potential of targeting MMP-14 in ACC. Using genetic silencing and pharmacologic inhibition of the hemopexin domain, we demonstrate that MMP-14 targeting reduces viability in both NCI-H295R ACC cells and patient-derived organoids (PTOs).

Further mechanistic investigation revealed that MMP-14 is a critical regulator of cell proliferation and DNA repair in ACC.

## RESULTS

### Elevated MMP-14 expression correlates with poor survival in ACC

We profiled the expression of 24 human MMP genes in ACC using the TCGA bulk RNA sequencing dataset. Of these, *MMP14* was the highest expressed MMP in ACC (**Fig. 1A**). Furthermore, We found that MMP14 expression increased with tumor stage (p=0.02, **Fig. 1B**), in contrast to most other cancers (**Fig. S1**), prompting further investigation into its functional role in ACC. Importantly, high *MMP-14* expression was significantly associated with worse overall survival (OS) (p = 0.0065, HR 5.01, 9% CI 1.56-16.14) and disease-free survival (DFS) (p=0.00035, HR 5.78, 95% CI 2.23-15.01) of ACC patients (**Figs. 1C - 1D**). These findings demonstrate that *MMP-14* is associated with ACC progression and represents a negative prognostic factor in ACC.

**Figure 1.**
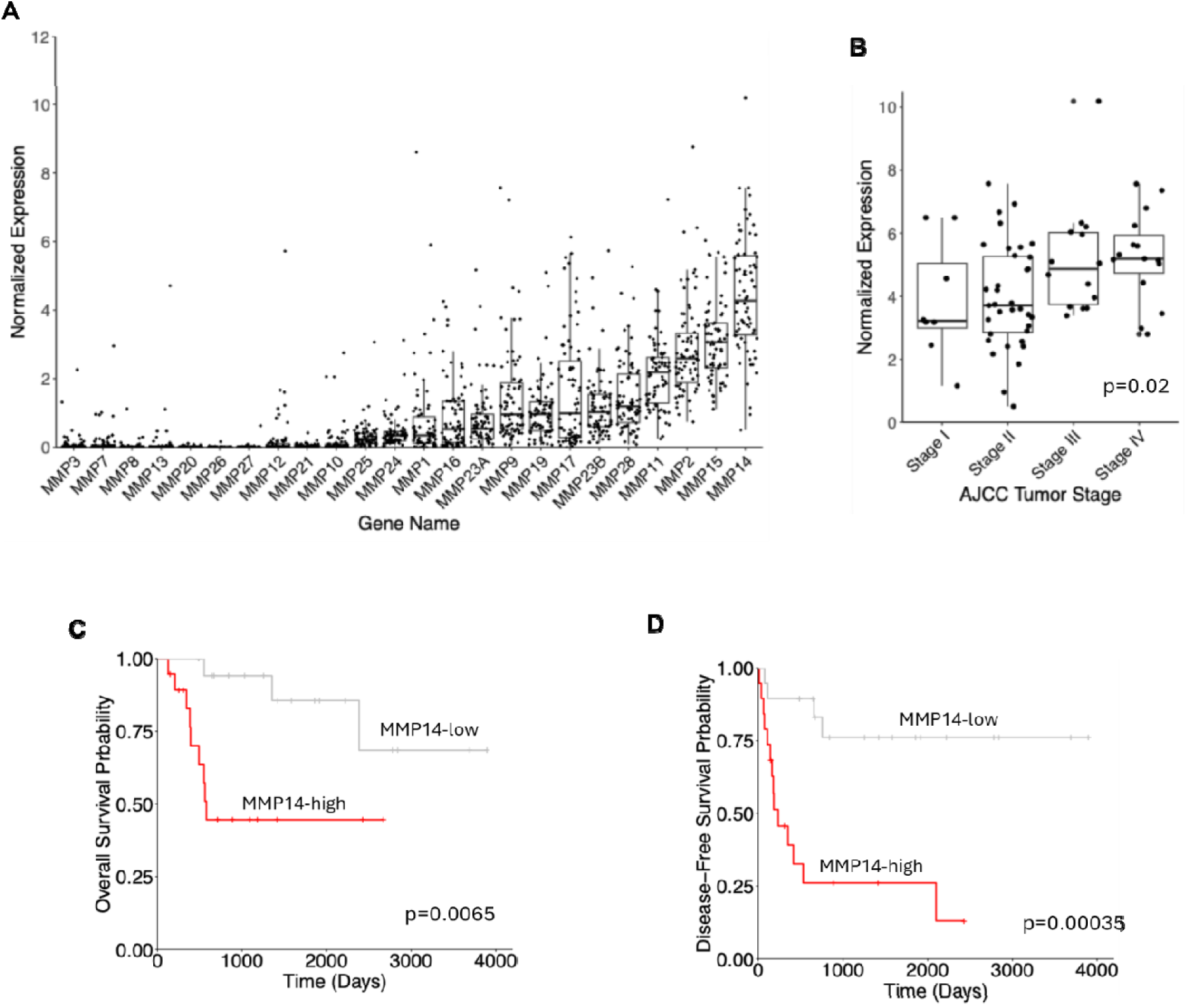
MMP-14 is upregulated and its elevation is associated with worse survival of ACC. **A**. MMP14 is the highest expressed MMPs in ACC. **B**. ANOVA trend analysis showing that increased expression of MMP14 is associated with advanced stage in ACC. **C** and **D**. Kaplan-Meier survival curves demonstrating that patients with high MMP14 expression exhibit decreased overall survival **(C**) and disease-free survival (**D**) compared to patients with low MMP14 expression.

### Inhibition of MMP-14 reduces ACC cell viability

#### NCI-H295R cell line

Because MMP14 is highly expressed in ACC and associated with poor outcomes, we investigated the functional role of MMP-14 using the ACC cell line, NCI-H295R. We first examined MMP-14 localization by immunofluorescence analysis, and revealed MMP-14 expression throughout cellular compartments, including strong perinuclear staining (**Figs. 2A and S2A**). This localization pattern suggests potential functions of MMP-14 beyond its canonical extracellular membrane-associated activities.

**Figure 2.**
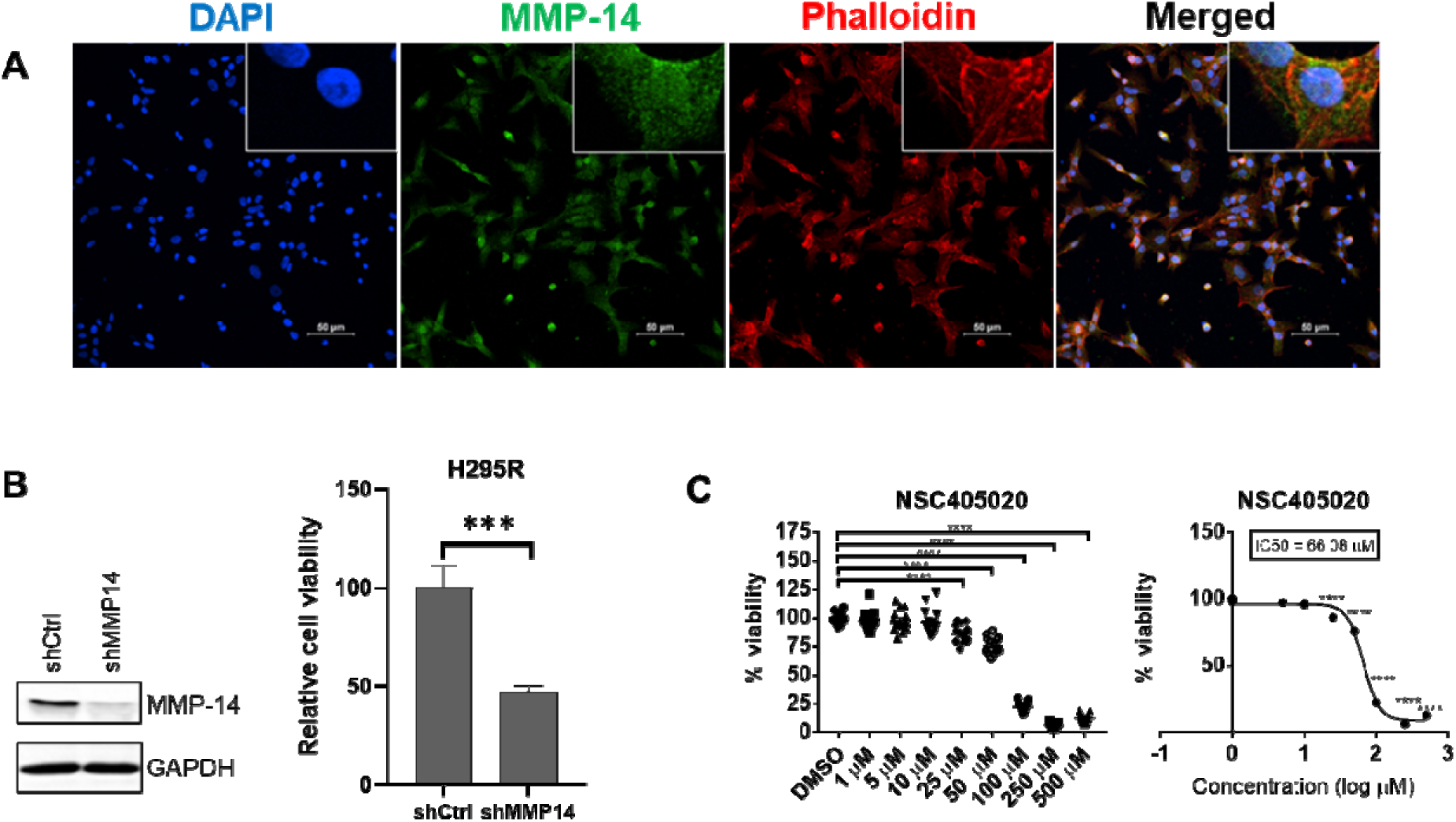
Genetic silencing or pharmacologic inhibition of MMP-14 reduces NCI-H295R cell viability. **A**. Immunofluorescence analysis of MMP-14 expression in NCI-H295R cells with Phalloidin staining to show actin. **B**. Genetic silencing MMP-14 by shRNA reduced NCI-H295R cell viability. NCI-H295R cells were transduced with lentivirus expressing control (shCtrl) or MMP-14 shRNA (shMMP14) for 72 hours and subjected to immunoblotting analysis of MMP-14 with GAPDH as loading control (Left panel), and cell viability analysis by CCK-8 (right panel). **C**. NCI-H295R cells were treated with increasing concentrations of MMP-14 inhibitor NSC405020 for 4 days and subjected to cell viability analysis. ns - not significant, * - p<0.05, ** - p < 0.01, *** - p<0.001, **** - p<0.0001.

We assessed MMP-14’s role in ACC cell survival using lentiviral short hairpin RNA (shRNA) to silence MMP-14 expression. Genetic knockdown of MMP-14 significantly reduced NCI-H295R cell viability (**Fig. 2B**), demonstrating that MMP-14 is essential for ACC cell survival. To examine its noncatalytic contributions[16, 25, 26] to this effect, we treated cells with NSC405020, a small-molecule inhibitor that specifically targets the hemopexin domain of MMP-14 without affecting catalytic activity[27]. Unlike traditional MMP inhibitors that target the catalytic domain, NSC405020 acts allosterically by binding the hemopexin domain and preventing homodimerization of MMP-14[27]. NSC405020 treatment reduced NCI-H295R cell viability in a dose-dependent manner (**Fig. 2C**). Together, these findings demonstrate that MMP-14 is essential for ACC cell growth and survival, in part due to its noncatalytic functions.

#### ACC patient-derived organoids (PTOs)

Patient-derived tumor organoids (PTOs) are 3D cultures that recapitulate key aspects of tumor biology *in vitro* and have emerged as powerful tools for preclinical therapeutic screening[28–31]. We previously established PTOs from ACC tumors that maintain adrenal cortex characteristics, including expression of the adrenal marker, SF-1 and steroid secretion[32]. To validate the role of MMP-14 in this physiologically relevant model system, we examined MMP-14 expression in three ACC PTO cultures (OSU1, OSU2, and OSU3). Immunofluorescence staining revealed MMP-14 expression across all three ACC PTOs, with relatively higher levels in OSU2 (**Figs. 3 and S2B**). We next tested whether MMP-14 inhibition affects PTO viability. Genetic silencing of MMP-14 significantly reduced ACC PTO viability (**Fig. 4A**), consistent with our observations in NCI-H295R cells. Similarly, NSC405020 treatment inhibited ACC PTO viability (**Fig. 4B**). These findings in both cell line and patient- derived organoid models demonstrate that MMP-14 represents a promising therapeutic target in ACC.

**Figure 3.**
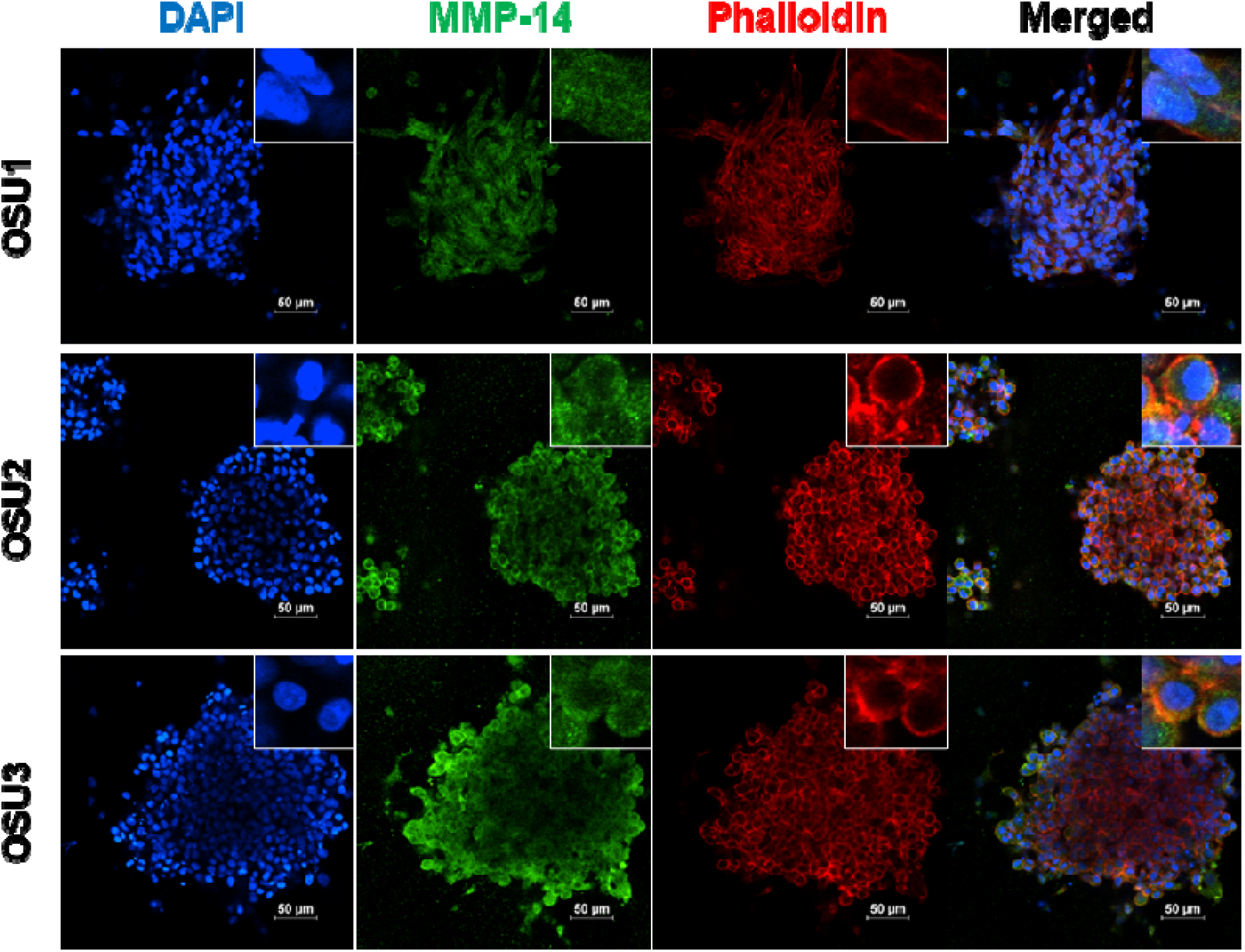
MMP-14 is expressed in ACC PTOs. Immunofluorescence demonstrating nuclei (DAPI, blue), MMP-14 (green), and actin (Phalloidin, red) in ACC PTO OSU1, OSU2 and OSU3.

**Figure 4.**
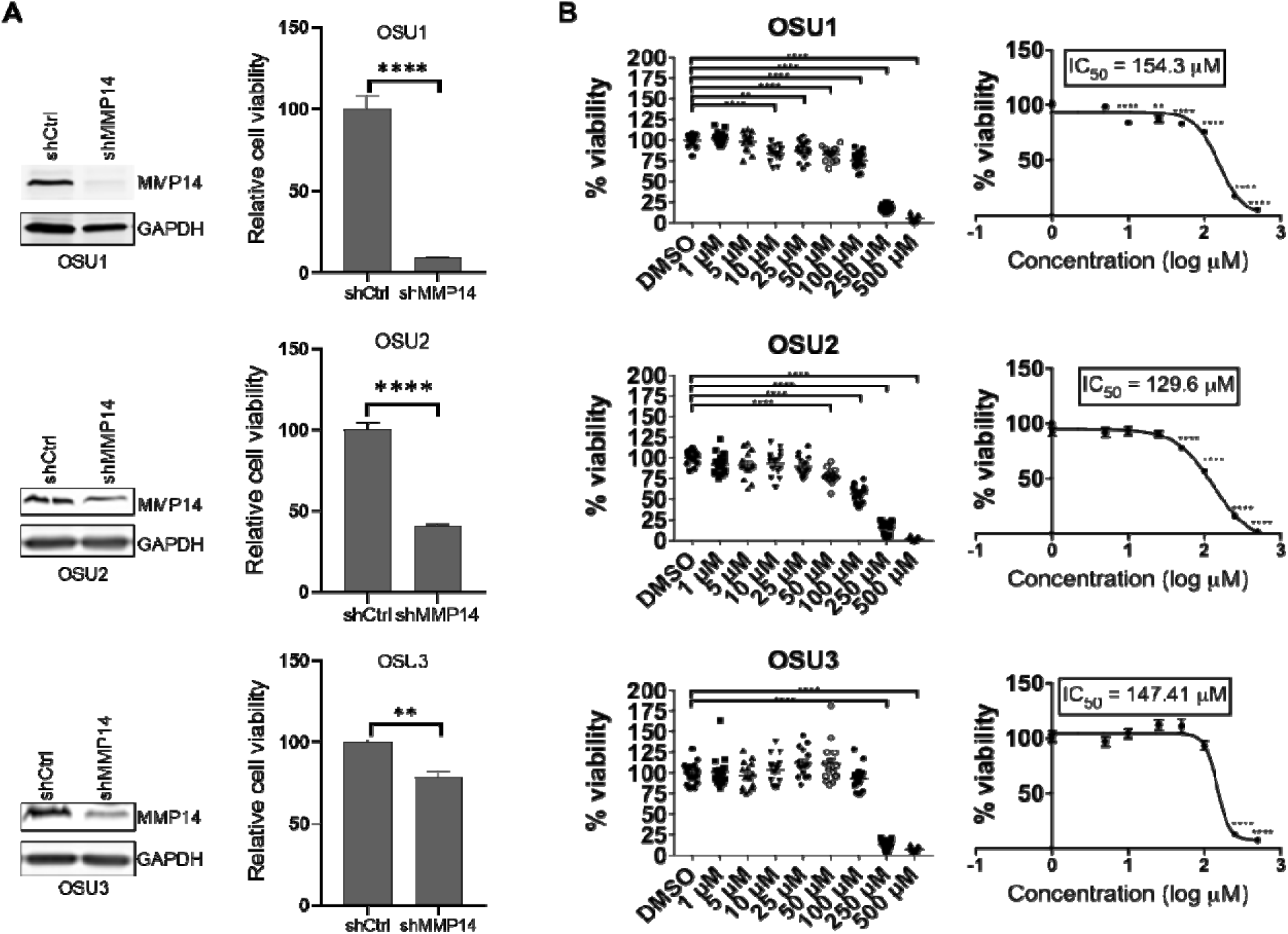
Genetic silencing or pharmacologic inhibition of MMP-14 reduces ACC PTOs viability. **A**. ACC PTO OSU1, OSU2 and OSU3 were transduced with lentivirus expressing control (shCtrl) or MMP-14 shRNA (shMMP14) for 72 hours and subjected to immunoblotting analysis of MMP-14 with GAPDH as loading control (Left panel), and cell viability analysis after 120 hours by CCK-8 (right panel). **B**. ACC PTO OSU1, OSU2 and OSU3 were treated with increasing concentrations of MMP-14 inhibitor NSC405020 for 120 hours and subjected to cell viability analysis. * - p<0.05, ** - p < 0.01, *** - p<0.001, **** - p<0.0001.

### MMP-14 downregulation leads to DNA replication checkpoint activation

To elucidate the mechanism by which MMP-14 maintains ACC cell viability, we investigated DNA damage response pathways. Prior studies in breast cancer cells demonstrated that MMP-14 modulates DNA damage response through stimulation of integrinβ1, which activates ERK and AKT signaling cascades[19, 33]. However, MMP-14 silencing in NCI-H295R cells did not alter AKT or ERK1/2 phosphorylation. Instead, we observed robust activation of CHK1 (**Fig. 5A**), a key kinase in the replication stress checkpoint pathway. CHK1 activation triggers downstream signaling that arrests cells in S-phase, providing time for DNA repair before replication continues[30]. To determine whether MMP-14 loss activates this checkpoint, we performed cell cycle analysis. Flow cytometry revealed that MMP-14 knockdown resulted in marked S-phase arrest in NCI-H295R cells (**Fig. 5B**), consistent with CHK1-mediated checkpoint activation.

**Figure 5.**
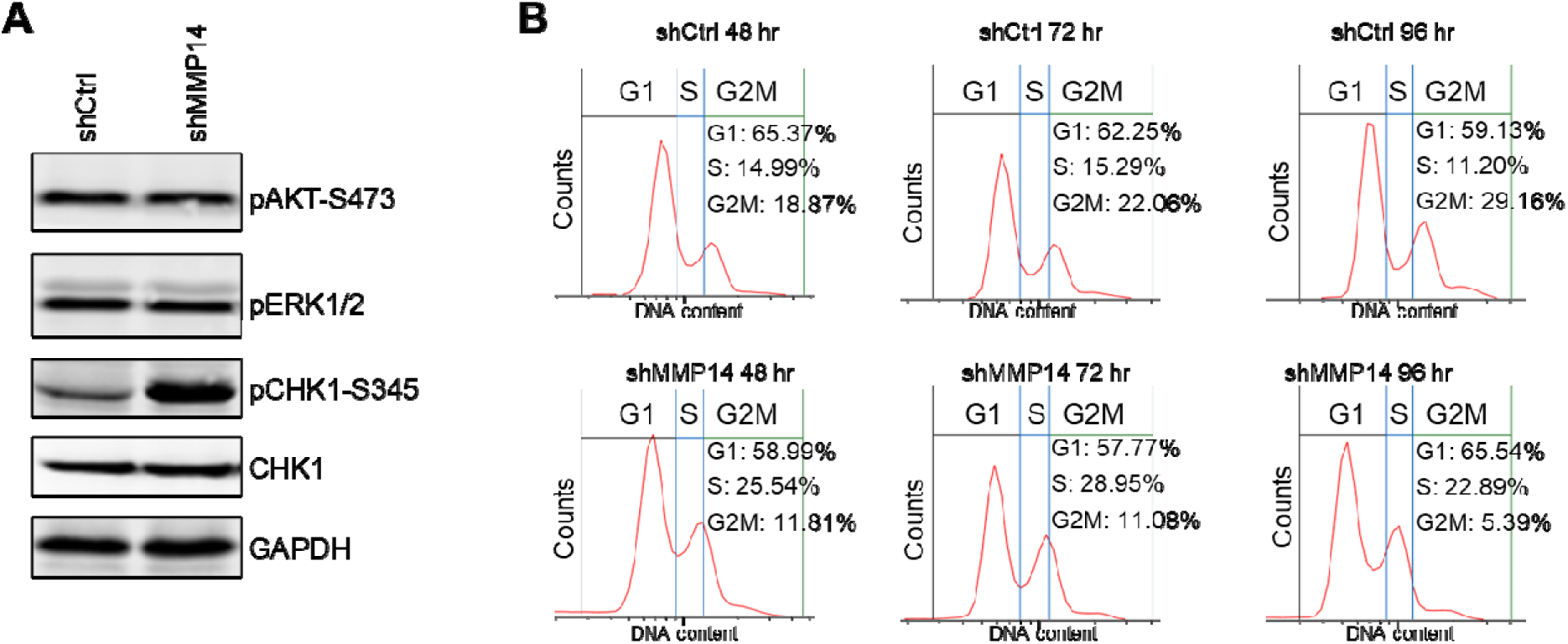
Genetic silencing of MMP-14 activates CHK1 checkpoint and induces S phase cell cycle arrest. **A**. NCI-H295R cells were transduced with lentivirus expressing control (shCtrl) or MMP-14 shRNA (shMMP14) for 72 hours and subjected to immunoblotting analysis of the indicated proteins with GAPDH as loading control. **B.** NCI-H295R cells were transduced with lentivirus expressing control (shCtrl) or MMP-14 shRNA (shMMP14) for 48, 72 and 96 hours. After staining DNA content by propidium iodine, the cells were subjected to cell cycle distribution analysis by flow cytometry.

### DNA damaging agents induce MMP-14 nuclear accumulation and chromatin binding

Another principal role of DNA damage response is to promote DNA damage repair. To investigate whether MMP-14 is functionally lined to DNA damage repair in ACC, we analyzed the TCGA dataset and found that higher MMP-14 expression correlated with elevated hallmark DNA repair signatures (**Fig. 6A**). Because nuclear translocation following DNA damage indicates recruitment to DNA repair pathways, we examined MMP-14 subcellular localization in response to DNA damage. We treated NCI-H295R cells with 4 Gy ionizing radiation (IR) and performed immunofluorescence analysis of MMP-14 at 1, 6, and 24 hours post-treatment. While MMP-14 was distributed throughout all cellular compartments under basal conditions, IR progressively induced MMP-14 nuclear accumulation over time (**Fig. 6B** and **Fig. S3**). Treatment with the DNA-damaging agent cisplatin produced similar results. Cisplatin (10 µM, 2 hours) increased nuclear accumulation of MMP-14 in NCI-H295R cells (**Fig. 6C** and **Fig. S4**). To further characterize MMP-14 redistribution following DNA damage, we performed subcellular fractionation after cisplatin treatment (10 µM, 2 and 6 hours), separating cytoplasmic, nuclear soluble, and chromatin-bound fractions. Cisplatin induced chromatin binding of the DNA damage response proteins TOPBP1 and RAD51, as expected. Notably, cisplatin also induced MMP-14 binding to chromatin (**Fig. 6D**), demonstrating that MMP-14 redistributes to chromatin in response to DNA damage.

**Figure 6.**
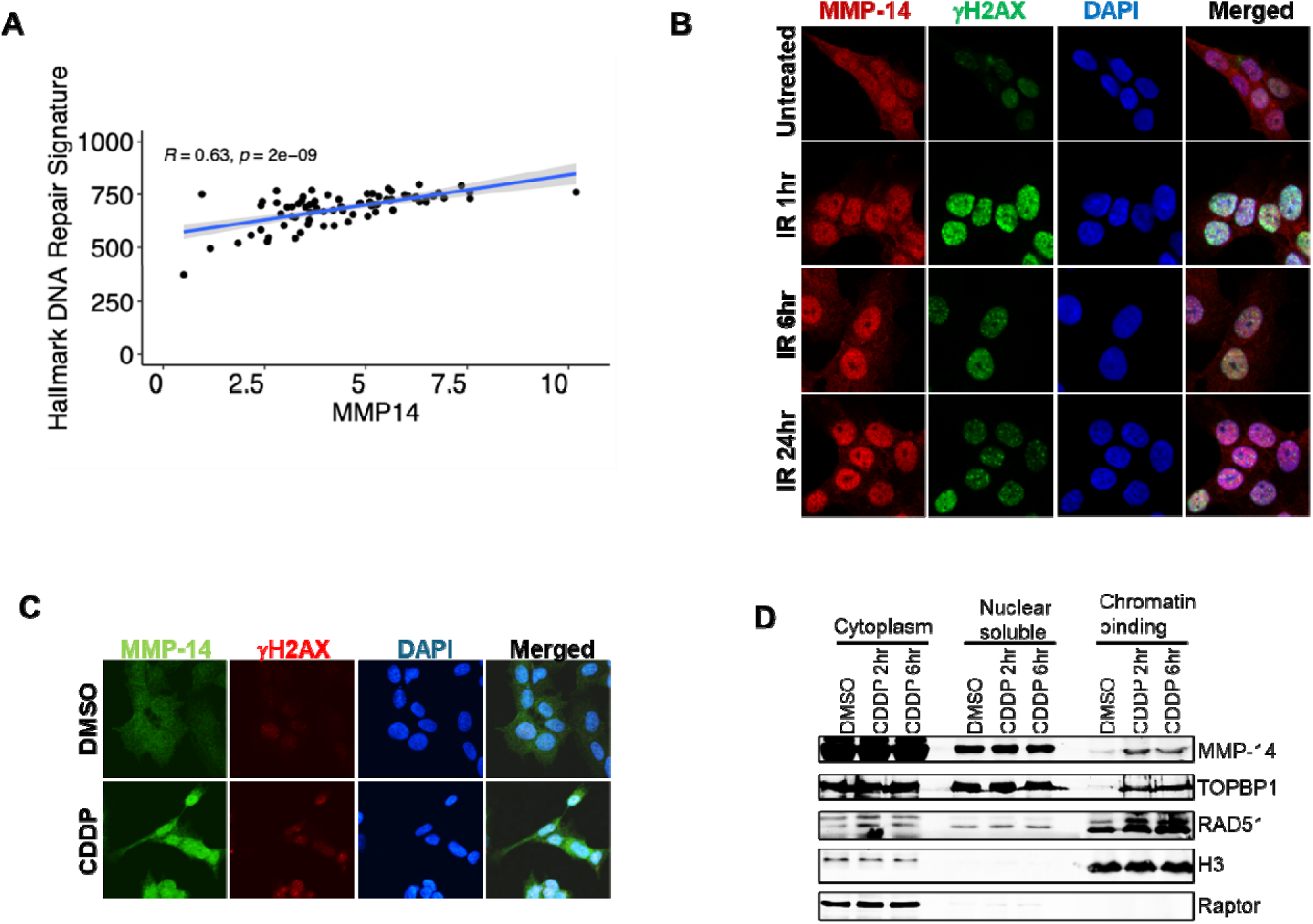
MMP-14 is associated with DNA damage response and repair. **A**. Increased *MMP14* expression is associated with increased hallmark DNA repair signature in the TCGA ACC dataset. **B**. Immunofluorescence analysis at 60X magnification of MMP-14 and γH2AX in NCI-H295R cells after treatment with 4 Gy X-ray for 1, 6 and 24 hours. **C**. Immunofluorescence analysis at 60X magnification of MMP-14 and γH2AX in NCI-H295R cells after treatment with 10 µM cisplatin (CDDP) for 2 hours. **D**. NCI-H295R cells were treated with or without cisplatin (CDDP, 10µM) for 2 and 6 hours. Cells were fractionated into cytoplasmic, nuclear soluble, and chromatin-bound fractions. Representative Western blots showing the levels of the indicated proteins in each fraction. H3 served as loading control for chromatin binding fraction; Raptor served as loading control for cytoplasm fraction.

### MMP-14 inhibition induces DNA damage and impairs DNA double strand break (DSBs) repair

The nuclear and chromatin localization of MMP-14 following DNA damage suggested a functional role in DNA repair. Consistent with this hypothesis, MMP-14 knockdown in NCI-H295R cells induced DNA damage, as demonstrated by increased γH2AX levels (**Fig. 7A**). Pharmacologic inhibition with NSC405020 produced the same effect (**Fig. 7B**). Similarly, both genetic and pharmacologic MMP-14 inhibition increased γH2AX levels in ACC PTOs (**Fig. 7C**-**7D**). The accumulation of γH2AX following MMP-14 inhibition indicates impaired repair of DNA double-strand breaks (DSBs). To identify which DSB repair pathway requires MMP-14 function, we employed homologous recombination (HR) and non-homologous end joining (NHEJ) reporter assays[34]. MMP-14 knockdown did not affect HR capacity, whereas

**Figure 7.**
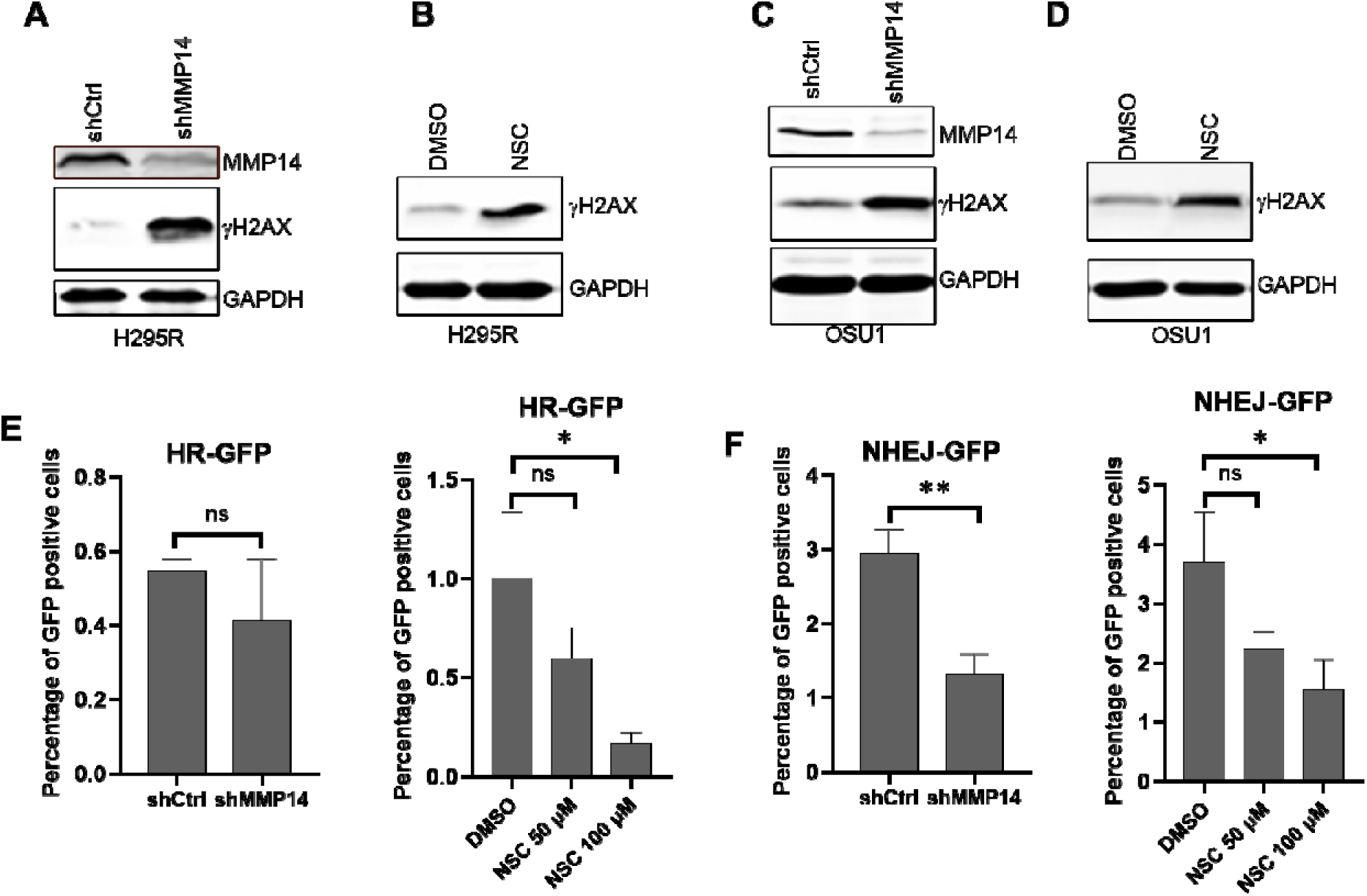
Genetic silencing or pharmacologic inhibition of MMP-14 induces DNA damages and reduces DSBs repair. **A**. NCI-H295R cells were transduced with lentivirus expressing control (shCtrl) or MMP-14 shRNA (shMMP14) for 72 hours and subjected to immunoblotting analysis of MMP-14 and γH2AX with GAPDH as loading control. **B**. NCI-H295R cells were treated with vehicle (DMSO) or 100 µM MMP-14 inhibitor NSC405020 for 24 hours and subjected to immunoblotting analysis of γH2AX with GAPDH as loading control. **C**. ACC PTO OSU1 were transduced with lentivirus expressing control (shCtrl) or MMP-14 shRNA (shMMP14) for 72 hours and subjected to immunoblotting analysis of γH2AX with GAPDH a loading control. **D**. ACC PTO OSU1 were treated with vehicle (DMSO) or 200 µM MMP-14 inhibitor NSC405020 for 48 hours and subjected to immunoblotting analysis of γH2AX with GAPDH as loading control. **E**. HR reporter assays of U2OS cells (stable reporter cell lines) after transduction of control or MMP14 shRNA for 48 hours (left panel) or treated with vehicle (DMSO) or MMP-14 inhibitor NSC405020 for 48 hours (right panel). **F**. NHEJ reporter assays of U2OS cells (stable reporter cell lines) after transduction of control or MMP14 shRNA for 48 hours (left panel) or treated with vehicle (DMSO) or MMP-14 inhibitor NSC405020 for 48 hours (right panel). ns - not significant, * - p<0.05, ** - p < 0.01.

NSC405020 treatment significantly reduced HR activity (**Fig. 7E**). In contrast, both genetic and pharmacologic MMP-14 inhibition substantially impaired NHEJ-mediated DSB repair (**Fig. 7F**). These data demonstrate that MMP-14 contributes to DSB repair primarily through the NHEJ pathway.

Collectively, these data demonstrate that MMP-14 maintains genome integrity during DNA replication through dual mechanisms: direct nuclear/chromatin association following DNA damage and CHK1-dependent checkpoint signaling, a pathway distinct from the integrinβ1/AKT/ERK signaling reported in breast cancer.

## DISCUSSION

Our study revealed that MMP-14 function extends beyond ECM remodeling to include non-canonical functions roles that contribute to ACC cell survival. In TCGA analysis, high expression of MMP-14, the most highly expressed MMP in ACC, was associated with worse overall and disease-free survival. Consistent with this clinical association, genetic and pharmacologic inhibition of MMP-14 reduced viability in NCI-H295R cells and PTOs. To elucidate the underlying mechanism, we investigated MMP-14’s intracellular functions and uncovered an unexpected role in genome maintenance: MMP-14 translocates to the nucleus following DNA damage and participates in NHEJ-mediated DNA repair.

The mechanism by which MMP-14 supports NHEJ remains to be elucidated. MMP-14 may interact with core NHEJ machinery, either through direct protein-protein interactions or by modulating chromatin accessibility at break sites. Alternatively, MMP-14 may regulate the activity or localization of NHEJ factors through proteolysis. Our data suggest that MMP-14 coordinates multiple aspects of genome maintenance through distinct structural elements. MMP-

14 knockdown induced robust CHK1 activation and S-phase checkpoint arrest. This response likely reflects the accumulation of unrepaired DNA damage and stalled replication forks when NHEJ is compromised. Interestingly, NSC405020 treatment, which targets the hemopexin domain without affecting catalytic activity, induced DNA damage but did not activate the CHK1/S-phase checkpoint (**Fig. S5**). These differential effects suggest that the ATR-CHK1 replication stress response is regulated by the MMP-14 catalytic domain, while NHEJ-mediated repair is mediated by the hemopexin domain.

Our findings reveal an intriguing paradox: while MMP-14 appears to maintain genome stability in ACC cells, prior studies in other cancer models demonstrate that MMP-14 promotes genomic instability[35]. MMP-14 localizes to centrosomes and cleaves pericentrin, a protein essential for mitotic spindle formation[36]. In mammary epithelial cells, MMP-14 overexpression disrupts centrosome function, resulting in abnormal spindle formation, aneuploidy, and chromosome instability[37], hallmarks of malignant transformation. Wali et al. further demonstrated that MMP-14 cleaves centrosomal BRCA2, regulating its removal from centrosomes during metaphase and disrupting centrosome duplication[38]. These destabilizing effects contrast with our observation that MMP-14 inhibition in ACC cells induces DNA damage, CHK1 activation, and impaired NHEJ repair.

Several explanations may reconcile this apparent contradiction. MMP-14 may exert opposing effects at different subcellular locations, with the dominant effect depending on cellular context. At centrosomes, MMP-14 cleavage of pericentrin and BRCA2 disrupts mitotic fidelity and promotes chromosomal instability, while in the nucleus, MMP-14 supports NHEJ-mediated repair. Cancer cells with high replication stress may depend on MMP-14’s nuclear repair function to tolerate ongoing DNA damage. In this context, loss of MMP-14 may trigger catastrophic DNA damage accumulation and checkpoint arrest. The tissue-specific nature of MMP-14 function is further evident when comparing ACC to breast cancer. In breast cancer cells, MMP-14 modulates DNA damage response through extracellular integrinβ1 signaling[19], whereas MMP-14 knockdown in ACC cells did not affect integrinβ1 downstream targets, as evidenced by unchanged AKT and ERK phosphorylation. The selective pressure in ACC—an aggressive cancer with limited treatment options—may have driven particular reliance on the direct DNA repair functions of MMP-14.

These considerations suggest that MMP-14 occupies a complex role in cancer biology, acting as both a driver of genomic instability during early transformation and a support for genome maintenance in established tumors. The observation that both genetic and pharmacologic MMP-14 inhibition reduces ACC PTO viability suggests therapeutic potential. NSC405020, which targets the hemopexin domain, impairs NHEJ and reduces cell viability, offering a more selective therapeutic approach than broad-spectrum MMP inhibitors. Importantly, targeting MMP-14 could sensitize ACC cells to DNA-damaging agents like cisplatin and ionizing radiation by impairing the repair of therapy-induced DNA damage.

Elucidating the molecular mechanisms by which MMP-14 contributes to NHEJ will be critical for developing targeted therapeutic strategies. Equally important will be understanding the relative contributions of centrosomal versus nuclear MMP-14 dysfunction to tumor cell lethality, which will inform whether inhibitors should target specific subcellular pools or global MMP-14 activity. Beyond these mechanistic insights, determining whether ACC’s dependency on MMP-14 represents a general feature of aggressive cancers or a tumor type-specific vulnerability will be essential for predicting which patients are most likely to benefit from MMP-14-targeted therapies.

This study establishes MMP-14 as a critical regulator of genome maintenance in ACC, extending well beyond its canonical role in ECM remodeling. Our finding that ACC cells depend on MMP14 for survival provides a mechanistic basis for the association between high MMP-14 expression and poor clinical outcomes. Targeting this novel dependency represents a promising therapeutic strategy that could sensitize ACC tumors to DNA-damaging chemotherapy and improve outcomes in this aggressive malignancy.

## MATERIALS AND METHODS

### NCI-H295R cell culture, chemicals and antibodies

The NCI-H295R ACC cell line was originally purchased from American Type Culture Collection (ATCC, Manassas, VA). Cells were maintained in pheno-red free DMEM/F12 with L-glutamine and 15 mM HEPES (Thermo Fisher Scientific, Waltham, MA) supplemented with 2.5% FBS (VWR, Radnor, PA), 1x ITS Premix Universal Cell Culture Supplement (Corning), and 1% penicillin-streptomycin (Millipore-Sigma, Burlington, MA). Cells were confirmed to be free of mycoplasma prior to experimental procedures. All cells utilized for experiments were under 25 passages in culture. MMP-14 inhibitor NSC405020 (cat.444295) was obtained from EMD Millipore (Burlington, MA). Anti-pERK1/2 (Thr202/Tyr204), ERK1/2, pAKT-S473, AKT, pCHK1-S345, CHK1, phospho-H2AX (Ser139), and GAPDH antibodies were purchased from Cell Signaling Technology (Danvers, MA).

### ACC tumor collection and PTO generation

ACC PTOs were generated as previously described[39]. Briefly, patients undergoing adrenalectomy for histology-confirmed ACC were consented for collection of specimens and clinical information using a protocol approved by The Ohio State University Institutional review board. A diagnosis of Long-lived ACC PTO cultures were generated from tissue specimens from three ACC patients with biochemical evidence of autonomous cortisol production (OSU1) or overt hypercortisolism (OSU2, OSU3). All PTO cultures were maintained in DMEM/F12 supplemented with 10 nM HEPES (Cytiva, Marlborough, MA), 1x Glutamax, 1x B27 Supplement (both Thermo Fisher Scientific), 1.25 mM n-acetylcysteine, 10 mM nicotinamide, 10 nM [Leu15]-Gastrin, 0.5 mM A-83-01, 1 mM PGE2 (all Millipore-Sigma), 0.5 mg/mL R-Spondin-1, 50 ng/mL EGF (both R&D Systems, Minneapolis, MN), 0.1 mg/mL FGF-10, 30 ng/mL recombinant murine Wnt3a, and 0.1 mg/mL recombinant human Noggin (all Thermo Fisher Scientific).

### Immunofluorescence

NCI-H295R cells were seeded on poly-L-lysine coated glass coverslips (Electron Microscopy Sciences, Hatfield, PA), fixed, and stained using the following antibodies: rabbit anti-human MMP-14 at 1:200 (Abcam, Cambridge, UK; #ab51074), phalloidin-Alexa Fluor^®^ 594 at 1:1000 (Abcam #ab176757), and goat anti-rabbit Alexa Fluor^®^ 488 at 1:1000 (Abcam #ab150081) or donkey anti-rabbit Alexa Fluor^®^ 647 at 1:1000 (Abcam #ab150063). NCI-H295R cells incubated in the absence of primary antibodies were used as negative controls. ACC PTOs were processed and stained for immunofluorescence microscopy using the following antibodies: rabbit anti-human MMP-14 at 1:200 (Abcam #ab51074), goat anti-rabbit Alexa Fluor^®^ 488 at 1:1000 (Abcam #ab150081), and phalloidin-Alexa Fluor^®^ 647 1:400 (Thermo Fisher Scientific #A2287). Negative control samples were stained with rabbit IgG isotype control at an equivalent concentration (Abcam #ab172730). All images were obtained by a Nikon Ti-2 Eclipse Confocal Fluorescence Microscope (Nikon, Tokyo, Japan).

### HR and NHEJ reporter assays

U2OS cells stably expressing fluorescent NHEJ (EJ5-GFP) and HR (DR-GFP) reporter assays were used as described previously[40, 41], and cultured in DMEM medium without sodium pyruvate (10-017-CV, Corning) supplemented with 10% FBS and 1% penicillin/streptomycin. Briefly, cells were transiently transfected with plasmids expressing I-SceI restriction enzyme for 24 hrs, followed by transduction with lentivirus expressing shCtrl or shMMP-14, or treated with different concentrations of NSC405020. After 48 hrs, GFP-positive cells were measured on a Cytek NL-3000 (CYTEK) flow cytometry.

### Cytotoxicity assays

#### NCI-H295R cytotoxicity assays

NCI-H295R cells were seeded at 1×10^4^ cells per well in a 96-well plate and allowed to adhere for 24 hours. Media was then replaced with fresh media containing specified drugs or equivalent DMSO vehicle control (Corning). Cell viability was assessed by water-soluble tertozolium-8 (WST-8) based Cell Counting Kit-8 assay (Dojindo, Mashiki, Kumamoto, Japan) per the manufacturer’s instructions at 4 days. Absorbance readings were conducted using a SpectraMax^®^ iD5 plate reader and SoftMax^®^ Pro Software (Molecular Devices, San Jose, CA). All assays were performed in biologic triplicates with 3-5 replicates per condition. The following drugs were used at concentrations ranging from 1 to 500000 nM: NSC405020. Mean viability of each drug treatment group was measured as a percentage of baseline DMSO vehicle treated cell absorbance. The half-maximal inhibitory concentration (IC_50_) for each drug was calculated by performing a four-parameter nonlinear best fit analysis of log concentration by percent viability on GraphPad Prism^®^ 7.0a software (GraphPad Software, La Jolla, CA). Differences in viability across each concentration of a particular drug were compared using one-factor ANOVA with post-hoc multiple comparison analysis by Dunnett’s test in GraphPad Prism^®^ 7.0a software (GraphPad Software). All data are reported as means ± standard error of the mean (SEM) with statistical significance set as p <0.05.

### ACC PTO cytotoxicity assays

Single-cell suspensions were generated by incubating well-established ACC organoid domes with 1 mg/mL dispase II (Thermo Fisher Scientific). 1.5×10^4^ cells per well were seeded in 96-well plates in 80% Matrigel^®^ as described above. Three days after seeding, media was replaced with media containing specified drugs or DMSO vehicle control. Organoid viability was assessed at 4 days of treatment by WST-8 assay as described above using the cell free Matrigel^®^ domes for background absorbance readings. Differences in viability across each concentration of a drug were compared using one-factor ANOVA with post-hoc multiple comparison analysis by Dunnett’s test in GraphPad Prism^®^ 7.0a software (GraphPad Software, La Jolla, CA). Each drug was evaluated in 5 replicates per condition across three or more independent experimental replicates.

### Gene silencing by lentivirus delivered shRNA

The plasmids of shRNAs of non-target control (SHC016VN) and targeting human MMP-14 (TRCN0000050855) were purchased from Sigma-Aldrich. Lentiviral particles for shRNA were produced in HEK293T cells transfected with shRNA plasmids, packaging (psPAX2) and envelope (pMD2G) plasmids (both from Addgene) using Lipofectamine 3000 (Invitrogen) according to manufacturer’s protocol. The lentiviral supernatants were harvested twice at 24- and 48-hour time points after transfection and passed through a membrane filter with 0.45 μm pore size. NCI-H295R cells were transduced with the lentiviral particles in the presence of 8 μg/mL polybrene (Sigma-Aldrich, #TR-1003-G). ACC PTOs were transduced in a similar manner, with minor modifications: ACC PTOs were dissociated with dispase prior to lentiviral transduction, and were plated in Matrigel 6 hours after transduction.

### Cell lysates preparation and immunoblotting

For NCI-H295R cells, cells cultured in plates were washed twice with ice-cold PBS, and lysed in RIPA lysis buffer (ThermoFisher) supplemented with 1x protease inhibitor (cOmplete, Roche) and phosphatase inhibitors (PhosSTOP, Roche) for 15 min on ice. Organoids cultured in Matrigel were mixed with 1 µg/mL dispase for 2 hours to completely resolve the Matrigel, followed by washing twice with ice-cold PBS, and lysed in RIPA lysis buffer supplemented with 1x protease inhibitor for 30 min on ice. The cell lysate was centrifuged at 150,000 rpm for 15 min at 4°C and the supernatant was collected. Protein quantification was determined with Dc protein assay kit (Bio-Rad, Hercules, CA). For immunoblotting, equal amounts of protein were resolved by SDS/PAGE (Bio-Rad) and transferred onto nitrocellulose membranes (cat. 1620115, Bio-Rad). Primary antibodies at a dilution of 1:1,000-5,000 were allowed to bind overnight at 4°C. After washing in Tris-buffered saline with 0.1% Tween 20 (TBS-Tween), membranes were incubated with a secondary antibody (1:5,000-10,000) conjugated to a near-infrared (NIR) fluorophore that binds to the primary antibody at room temperature for 1 hour in the dark. Membranes were washed with TBS-Tween and scanned using an Azure imaging system (Azure Biosystem) to detect the fluorescence signal. Secondary antibodies of donkey anti-rabbit IRDye 680RD (926-68073), donkey anti-rabbit IRDye 800CW (926-32212), and donkey anti-mouse IRDye 680RD (926-68072) were purchased from LI-COR.

### Isolation of subcellular fractions and chromatin-bound proteins

To isolate cytoplasmic, soluble nuclear, and chromatin-binding proteins, the cells were lysed in 500 µl ice-cold Buffer A [50 mM HEPES-KOH (pH 7.5), 140 mM NaCl, 1 mM EDTA (pH 8.0), 10% glycerol, 0.5% Nonidet P-40, 0.25% Triton X-100, 1 mM DTT, 1×protease inhibitor cocktail]. After centrifugation (700×g, 10 min), the supernatant was collected as the cytoplasm fraction and pellets were washed with Buffer A and resuspended in ice-cold Buffer B [10 mM Tris–HCl (pH 8.0), 200 mM NaCl, 1mM EDTA (pH 8.0), 0.5 mM EGTA (pH 8.0)], 1×protease inhibitor cocktail]. After extraction and centrifugation (20,000×g, 10 min), the soluble nuclear extract (NE) was collected. The pellet was washed with Buffer B and resuspended in Buffer C [500 mM Tris-HCl (pH 6.8), 500 mM NaCl, 1×protease inhibitor cocktail]. The suspension was sonicated for 2×10s and centrifuged (20,000×g, 10 min), and the supernatant was collected for chromatin binding fraction.

### Flow cytometric analysis for cell cycle phases

Around 0.2-5 x 10^6^ NCI-H295R cells were seeded per well of a 12-well plate and incubated for 24 hours. Cells were treated with the specified compounds for the specified duration. All sample wells were harvested by trypsinization. Cell pellets were washed with PBS and were then then fixed with 70% ethanol at 4°C overnight. All samples were then incubated with 10 µg/mL propidium iodide and 200 μg/mL RNAse A for 1-2 hours at 37 °C. Samples were then stored at 4 °C until analyzed on a Cytek NL-3000 flow cytometry within 1-2 days.

### TCGA data analysis

TCGA RNA sequencing dataset containing log2(TPM+1) normalized data was downloaded from the UCSC Treehouse Tumor Compendium v11 Public PolyA cohort. The dataset was filtered to include adult ACC patients only (age at diagnosis >= 18 years old); as a result, gene expression data for 74 ACC patients were analyzed. *ggplot2* v3.5 for R v4.1 [42] was used to compare log2(TPM+1) expression values of all human MMP genes in ACC. ANOVA trend analysis was performed in R v.4.1. For the pan-cancer ANOVA trend analysis, 21 cancer types were included. Hematological cancers, glioma, glioblastoma, and cancers with fewer than 10 patients were excluded from analysis. To perform survival analysis, packages *survival* v. 3.2 and *survminer* v. 0.4 were used[43] (https://rdrr.io/cran/survminer/). For the overall and the disease-free survival analyses, ACC patients with *MMP14* expression in the upper quartile (n = 19) were considered to have high *MMP14* expression, whereas ACC patients with *MMP14* expression in the lower quartile (n = 19) were considered to have low *MMP14* expression. The Hallmark DNA repair signature was downloaded from the Molecular Signatures Database (MSigDB)[44].

### Statistical analysis

Graphs for cytotoxicity assays were constructed using GraphPad Prism (GraphPad Software, San Diego, CA). All data are presented as mean ± SEM. Log-rank tests were used to calculate P-values in Kaplan–Meier analysis. Wilcoxon mean rank-sum tests were used for two-group comparison of continuous variables. Statistical significance was determined by unpaired two-tailed t-test or two-way ANOVAs. P< 0.05 was considered statistically significant.

## Acknowledgements

We acknowledge resources from the Campus Microscopy and Imaging Facility, Biospecimen Services Shared Resource, Flow Cytometry Shared Resource, Immune Monitoring and Discovery Platform, and the Microscopy Shared Resource. These shared resources are supported in part by the Cancer Center Support Grant P30 CA016058, National Cancer Institute, Bethesda, MD. We thank Dr. Jeremy Stark (City of Hope, Duarte, CA) for providing NHEJ (EJ5-GFP) and HR (DR-GFP) reporter system. We also acknowledge Matthew Ringel, MD, Wayne Miles, PhD, Abby Green, MD, PhD, Matthew Summers, PhD, and Monica Venere, PhD for insightful discussions.

## Author contributions

Conceptualization: P.H.D. Methodology: C.S. L.V.P. and P.H.D. Investigation: C.S., D.M.C, L.V.P., Y.S., C.A., Z.L., S.P., V.T., J.E.P, B.S.M., B.B. and H.L.

Specimen Acquisition: P.H.D., J.E.P., and B.S.M. Formal analysis: C.S., L.V.P. and D.M.C.

Funding acquisition: P.H.D. Project administration, supervision, and validation: P.H.D. Writing-original draft: C.S., L.V.P., D.M.C. Writing-review and editing: C.S., L.V.P., and P.H.D.

---

**Figure S1.**
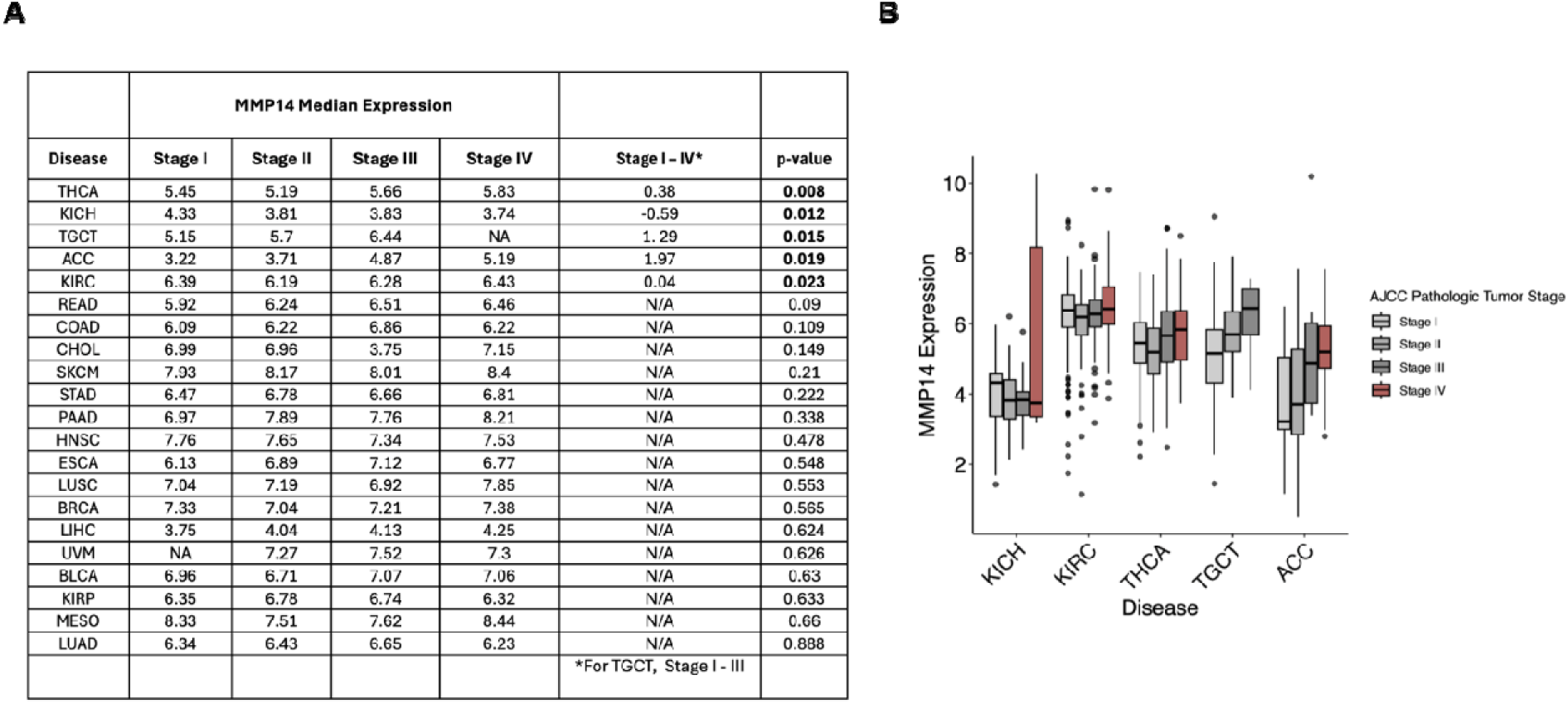
Pan-Cancer ANOVA Trend Analysis. A. The table demonstrating median MMP14 expression values for stage I, stage II, stage III, and stage IV cancers, stage I-IV slope, and ANOVA trend analysis p-values. B. Boxplots representing MMP14 expression in stage I-IV cancers associated with significant ANOVA trend analysis p-values.

**Figure S2.**
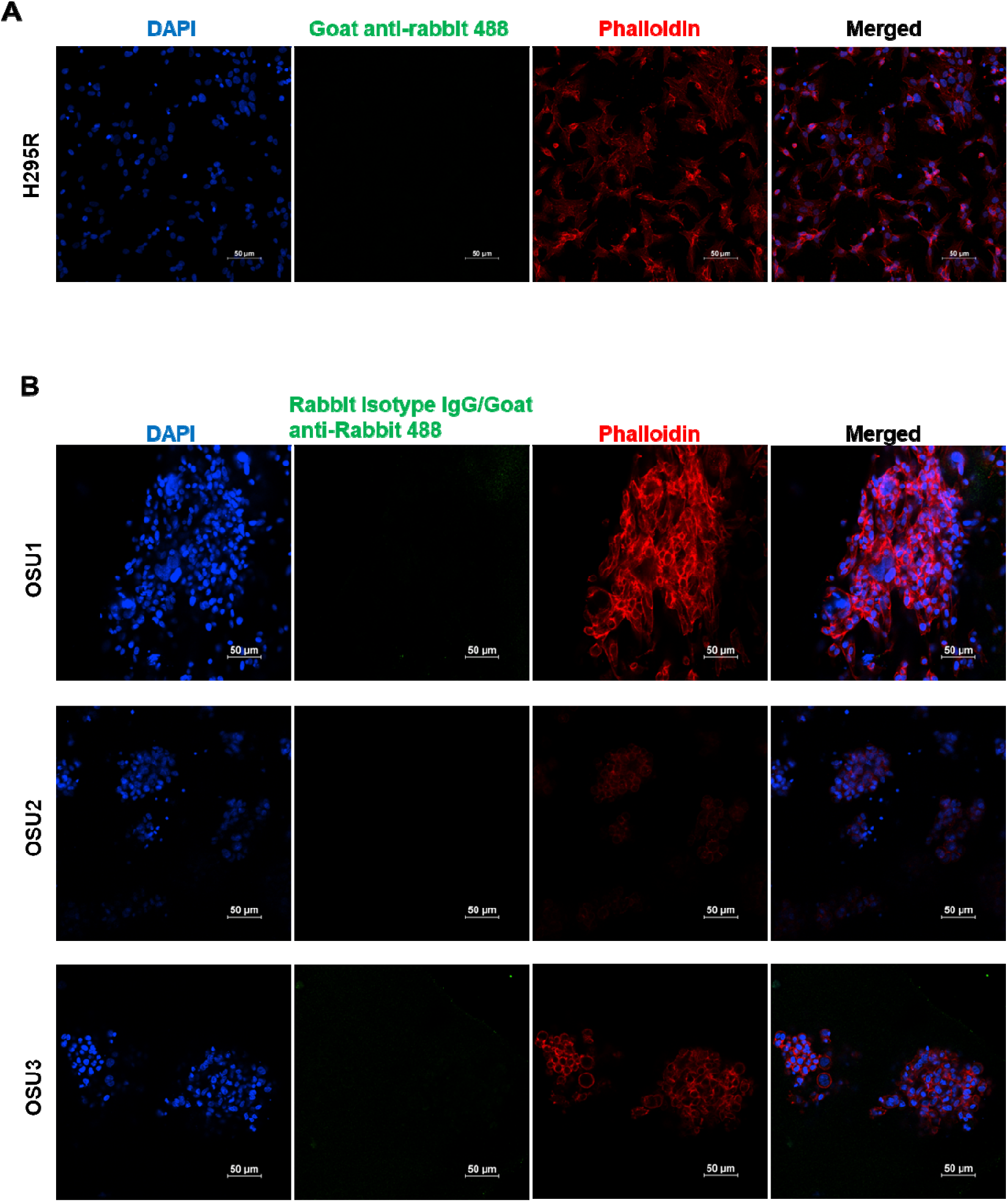
Isotype antibody controls for MMP-14 immunofluorescence. **A**. Isotype antibody controls for MMP-14 immunofluorescence in NCI-H295R cells. Related to Figure 2A. **B**. Isotype antibody controls for MMP-14 immunofluorescence in ACC PTOs. Related to Figure 3.

**Figure S3.**
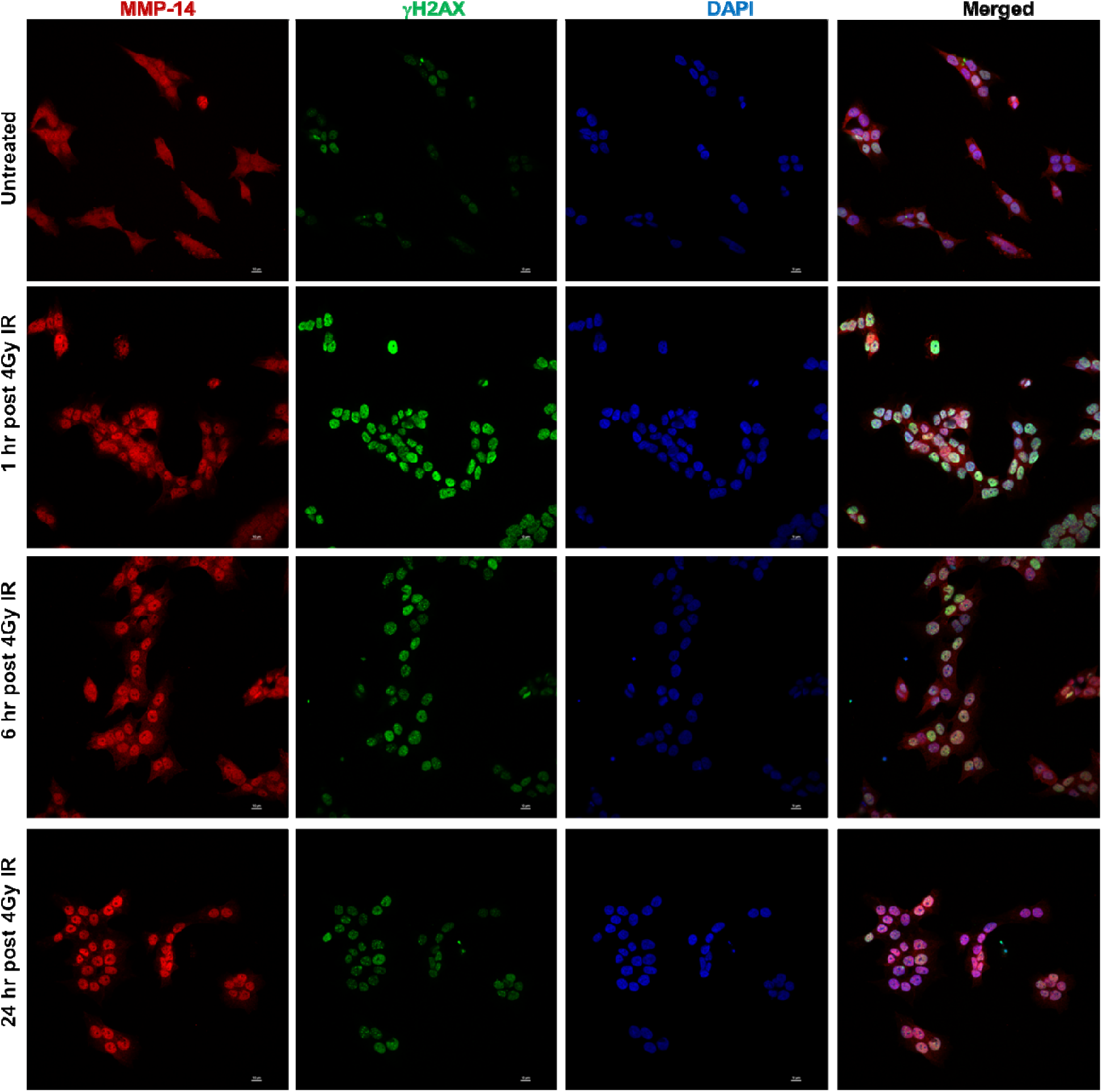
MMP-14 undergoes nuclear accumulation in response to ionizing radiation. Related to Figure 6B.

**Figure S4.**
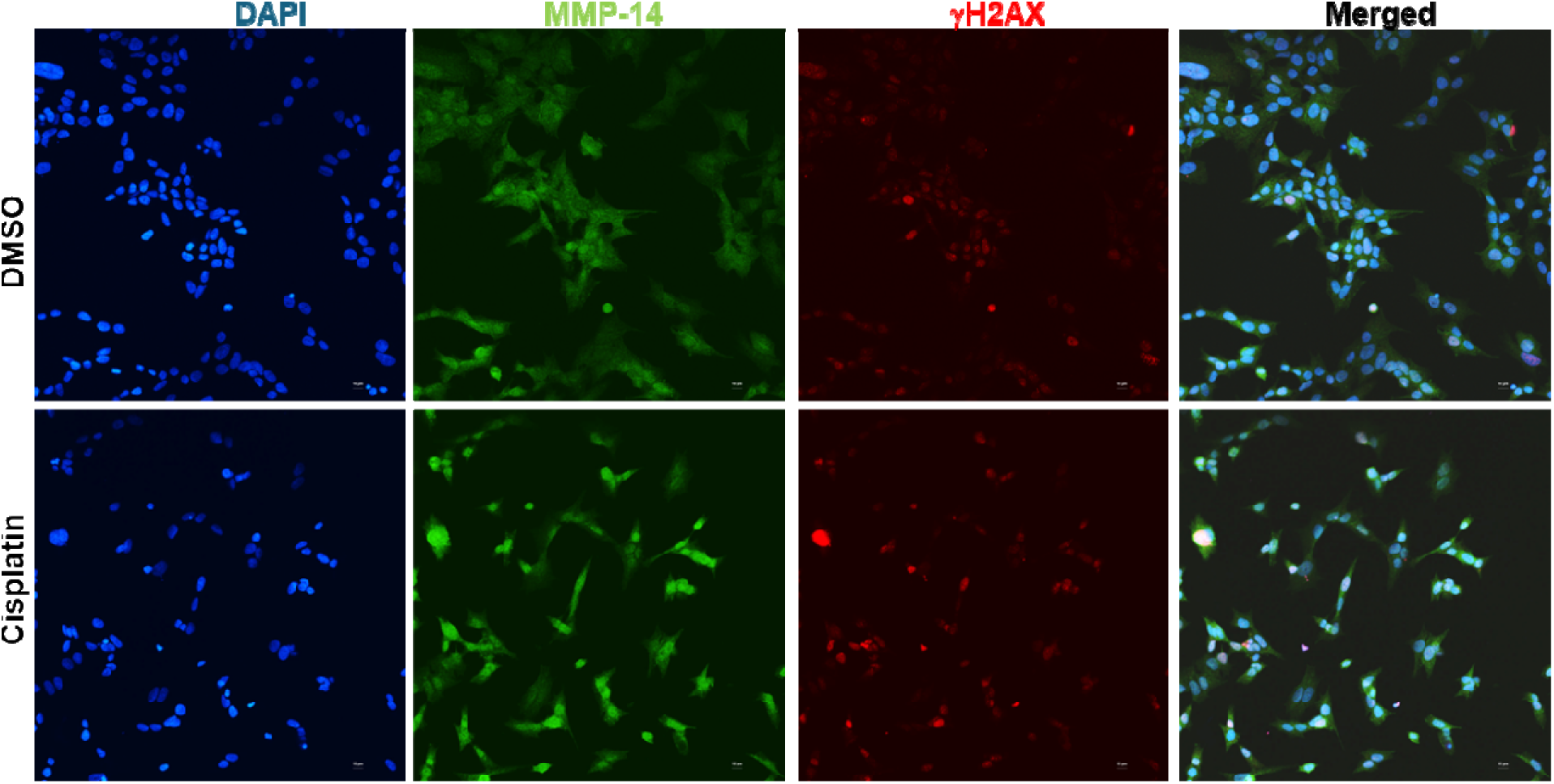
MMP-14 undergoes nuclear accumulation in response to cisplatin treatment. Related to Figure 6C.

**Figure S5.**
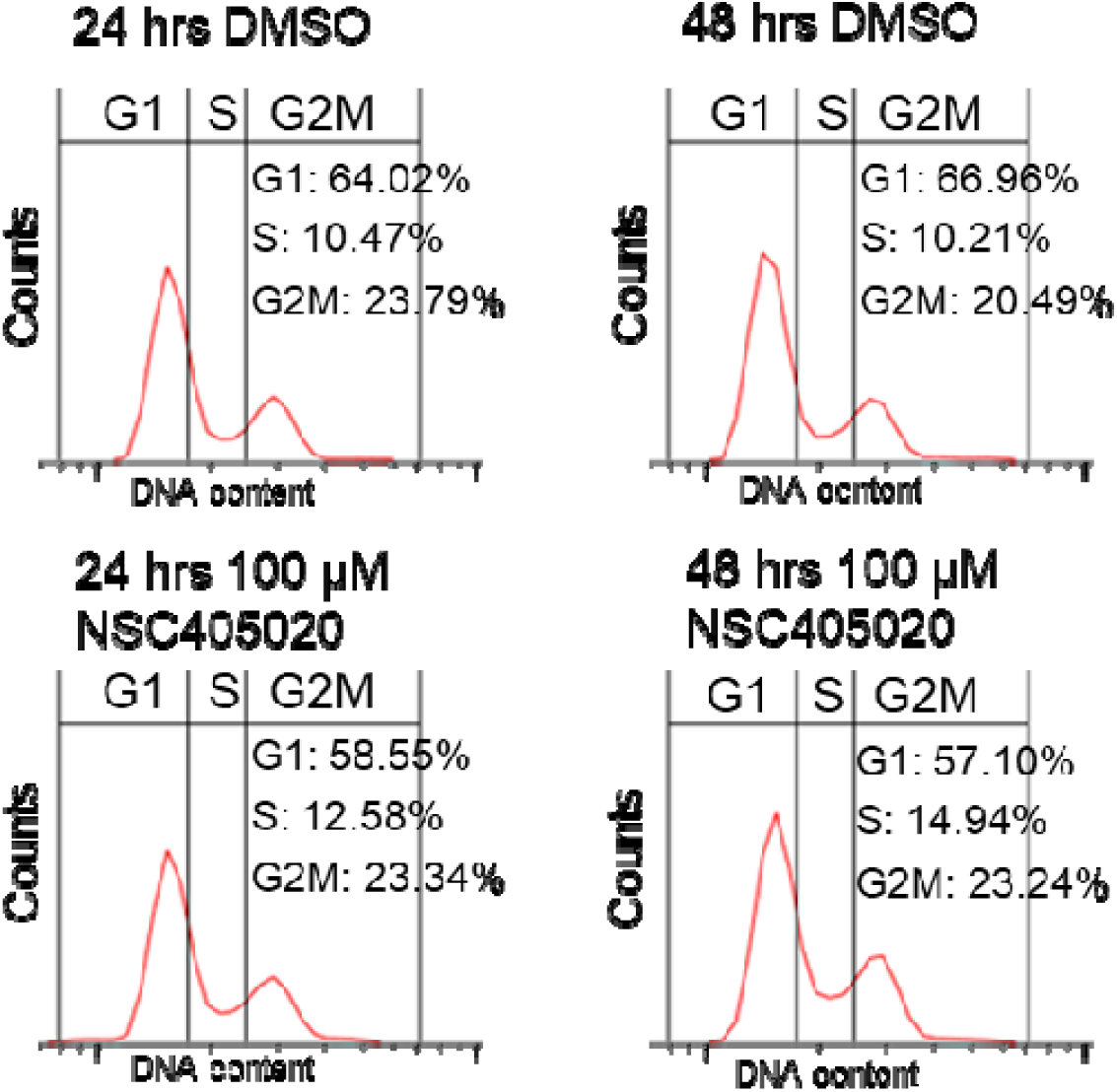
NSC405020 inhibition of MMP-14 does not induce S-phase cell cycle arrest. Related to Figure 5.

## REFERENCES

1. Costa R, Carneiro BA, Tavora F, Pai SG, Kaplan JB, Chae YK, et al. The challenge of developmental therapeutics for adrenocortical carcinoma. Oncotarget. 2016;7:46734–49. 10.18632/oncotarget.8774.

2. Fassnacht M, Terzolo M, Allolio B, Baudin E, Haak H, Berruti A, et al. Combination chemotherapy in advanced adrenocortical carcinoma. N Engl J Med. 2012;366:2189–97. 10.1056/NEJMoa1200966.

3. Schmid P, Sablin M-P, Bergh J, Im S-A, Lu Y-S, Martínez N, et al. A phase Ib/II study of xentuzumab, an IGF-neutralising antibody, combined with exemestane and everolimus in hormone receptor-positive, HER2-negative locally advanced/metastatic breast cancer. Breast Cancer Res BCR. 2021;23:8. 10.1186/s13058-020-01382-8.

4. Naing A, Lorusso P, Fu S, Hong D, Chen HX, Doyle LA, et al. Insulin growth factor receptor (IGF-1R) antibody cixutumumab combined with the mTOR inhibitor temsirolimus in patients with metastatic adrenocortical carcinoma. Br J Cancer. 2013;108:826–30. 10.1038/bjc.2013.46.

5. Miller KC, Chintakuntlawar AV, Hilger C, Bancos I, Morris JC, Ryder M, et al. Salvage Therapy With Multikinase Inhibitors and Immunotherapy in Advanced Adrenal Cortical Carcinoma. J Endocr Soc. 2020;4:bvaa069. 10.1210/jendso/bvaa069.

6. Fassnacht M, Puglisi S, Kimpel O, Terzolo M. Adrenocortical carcinoma: a practical guide for clinicians. Lancet Diabetes Endocrinol. 2025;13:438–52. 10.1016/S2213-8587(24)00378-4.

7. Kessenbrock K, Plaks V, Werb Z. Matrix metalloproteinases: regulators of the tumor microenvironment. Cell. 2010;141:52–67. 10.1016/j.cell.2010.03.015.

8. Koshikawa N, Schenk S, Moeckel G, Sharabi A, Miyazaki K, Gardner H, et al. Proteolytic processing of laminin-5 by MT1-MMP in tissues and its effects on epithelial cell morphology. FASEB J Off Publ Fed Am Soc Exp Biol. 2004;18:364–6. 10.1096/fj.03-0584fje.

9. Zarrabi K, Dufour A, Li J, Kuscu C, Pulkoski-Gross A, Zhi J, et al. Inhibition of matrix metalloproteinase 14 (MMP-14)-mediated cancer cell migration. J Biol Chem. 2011;286:33167–77. 10.1074/jbc.M111.256644.

10. Stawowczyk M, Wellenstein MD, Lee SB, Yomtoubian S, Durrans A, Choi H, et al. Matrix Metalloproteinase 14 promotes lung cancer by cleavage of Heparin-Binding EGF-like Growth Factor. Neoplasia N Y N. 2017;19:55–64. 10.1016/j.neo.2016.11.005.

11. Niland S, Riscanevo AX, Eble JA. Matrix Metalloproteinases Shape the Tumor Microenvironment in Cancer Progression. Int J Mol Sci. 2021;23:146. 10.3390/ijms23010146.

12. Shimoda M, Khokha R. Metalloproteinases in extracellular vesicles. Biochim Biophys Acta Mol Cell Res. 2017;1864 11 Pt A:1989–2000. 10.1016/j.bbamcr.2017.05.027.

13. Koziol A, Gonzalo P, Mota A, Pollán Á, Lorenzo C, Colomé N, et al. The protease MT1-MMP drives a combinatorial proteolytic program in activated endothelial cells. FASEB J Off Publ Fed Am Soc Exp Biol. 2012;26:4481–94. 10.1096/fj.12-205906.

14. Xia X-D, Alabi A, Wang M, Gu H-M, Yang RZ, Wang G-Q, et al. Membrane-type I matrix metalloproteinase (MT1-MMP), lipid metabolism, and therapeutic implications. J Mol Cell Biol. 2021;13:513–26. 10.1093/jmcb/mjab048.

15. Yang J, Kasberg WC, Celo A, Liang Z, Quispe K, Stack MS. Post-translational modification of the membrane type 1 matrix metalloproteinase (MT1-MMP) cytoplasmic tail impacts ovarian cancer multicellular aggregate dynamics. J Biol Chem. 2017;292:13111–21. 10.1074/jbc.M117.800904.

16. D’Alessio S, Ferrari G, Cinnante K, Scheerer W, Galloway AC, Roses DF, et al. Tissue Inhibitor of Metalloproteinases-2 Binding to Membrane-type 1 Matrix Metalloproteinase Induces MAPK Activation and Cell Growth by a Non-proteolytic Mechanism. J Biol Chem. 2008;283:87–99. 10.1074/jbc.M705492200.

17. Valacca C, Tassone E, Mignatti P. TIMP-2 Interaction with MT1-MMP Activates the AKT Pathway and Protects Tumor Cells from Apoptosis. PLOS ONE. 2015;10:e0136797. 10.1371/journal.pone.0136797.

18. Sakamoto T, Seiki M. Cytoplasmic tail of MT1-MMP regulates macrophage motility independently from its protease activity. Genes Cells Devoted Mol Cell Mech. 2009;14:617–26. 10.1111/j.1365-2443.2009.01293.x.

19. Thakur V, Zhang K, Savadelis A, Zmina P, Aguila B, Welford SM, et al. The membrane tethered matrix metalloproteinase MT1-MMP triggers an outside-in DNA damage response that impacts chemo- and radiotherapy responses of breast cancer. Cancer Lett. 2019;443:115–24. 10.1016/j.canlet.2018.11.031.

20. López-Otín C, Overall CM. Protease degradomics: a new challenge for proteomics. Nat Rev Mol Cell Biol. 2002;3:509–19. 10.1038/nrm858.

21. Chen W, Huang S, Shi K, Yi L, Liu Y, Liu W. Prognostic Role of Matrix Metalloproteinases in Cervical Cancer: A Meta-Analysis. Cancer Control J Moffitt Cancer Cent. 2021;28:10732748211033743. 10.1177/10732748211033743.

22. Zhang L, Jin S, Wei Y, Wang C, Zou H, Hu J, et al. Prognostic Significance of Matrix Metalloproteinase 14 in Patients with Cancer: a Systematic Review and Meta-Analysis. Clin Lab. 2020;66. 10.7754/Clin.Lab.2019.190831.

23. Liu J, Li Y, Lian X, Zhang C, Feng J, Tao H, et al. Potential target within the tumor microenvironment - MT1-MMP. Front Immunol. 2025;16:1517519. 10.3389/fimmu.2025.1517519.

24. Li M, Li S, Zhou L, Yang L, Wu X, Tang B, et al. Immune Infiltration of MMP14 in Pan Cancer and Its Prognostic Effect on Tumors. Front Oncol. 2021;11:717606. 10.3389/fonc.2021.717606.

25. Gifford V, Itoh Y. MT1-MMP-dependent cell migration: proteolytic and non-proteolytic mechanisms. Biochem Soc Trans. 2019;47:811–26. 10.1042/BST20180363.

26. Attur M, Lu C, Zhang X, Han T, Alexandre C, Valacca C, et al. Membrane-type 1 Matrix Metalloproteinase Modulates Tissue Homeostasis by a Non-proteolytic Mechanism. iScience. 2020;23:101789. 10.1016/j.isci.2020.101789.

27. Remacle AG, Golubkov VS, Shiryaev SA, Dahl R, Stebbins JL, Chernov AV, et al. Novel MT1-MMP small-molecule inhibitors based on insights into hemopexin domain function in tumor growth. Cancer Res. 2012;72:2339–49. 10.1158/0008-5472.CAN-11-4149.

28. Tuveson D, Clevers H. Cancer modeling meets human organoid technology. Science. 2019;364:952–5. 10.1126/science.aaw6985.

29. Dedhia PH, Bertaux-Skeirik N, Zavros Y, Spence JR. Organoid Models of Human Gastrointestinal Development and Disease. Gastroenterology. 2016;150:1098–112. 10.1053/j.gastro.2015.12.042.

30. Skardal A, Sivakumar H, Rodriguez MA, Popova LV, Dedhia PH. Bioengineered in vitro three-dimensional tumor models in endocrine cancers. Endocr Relat Cancer. 2024;31:e230344. 10.1530/ERC-23-0344.

31. Drost J, Clevers H. Organoids in cancer research. Nat Rev Cancer. 2018;18:407–18. 10.1038/s41568-018-0007-6.

32. Sbiera S, Schmull S, Assie G, Voelker H-U, Kraus L, Beyer M, et al. High diagnostic and prognostic value of steroidogenic factor-1 expression in adrenal tumors. J Clin Endocrinol Metab. 2010;95:E161–171. 10.1210/jc.2010-0653.

33. Thakur V, Tiburcio de Freitas J, Li Y, Zhang K, Savadelis A, Bedogni B. MT1-MMP-dependent ECM processing regulates laminB1 stability and mediates replication fork restart. PloS One. 2021;16:e0253062. 10.1371/journal.pone.0253062.

34. Wright WD, Shah SS, Heyer W-D. Homologous recombination and the repair of DNA double-strand breaks. J Biol Chem. 2018;293:10524–35. 10.1074/jbc.TM118.000372.

35. Golubkov VS, Boyd S, Savinov AY, Chekanov AV, Osterman AL, Remacle A, et al. Membrane type-1 matrix metalloproteinase (MT1-MMP) exhibits an important intracellular cleavage function and causes chromosome instability. J Biol Chem. 2005;280:25079–86. 10.1074/jbc.M502779200.

36. Golubkov VS, Chekanov AV, Doxsey SJ, Strongin AY. Centrosomal pericentrin is a direct cleavage target of membrane type-1 matrix metalloproteinase in humans but not in mice: potential implications for tumorigenesis. J Biol Chem. 2005;280:42237–41. 10.1074/jbc.M510139200.

37. Golubkov VS, Chekanov AV, Savinov AY, Rozanov DV, Golubkova NV, Strongin AY. Membrane type-1 matrix metalloproteinase confers aneuploidy and tumorigenicity on mammary epithelial cells. Cancer Res. 2006;66:10460–5. 10.1158/0008-5472.CAN-06-2997.

38. Wali N, Hosokawa K, Malik S, Saito H, Miyaguchi K, Imajoh-Ohmi S, et al. Centrosomal BRCA2 is a target protein of membrane type-1 matrix metalloproteinase (MT1-MMP). Biochem Biophys Res Commun. 2014;443:1148–54. 10.1016/j.bbrc.2013.12.103.

39. Popova LV, Garfinkle EAR, Chopyk DM, Navarro JB, Rivaldi A, Shu Y, et al. Single Nuclei Sequencing Reveals Intratumoral Cellular Heterogeneity and Replication Stress in Adrenocortical Carcinoma. BioRxiv Prepr Serv Biol. 2024;:2024.09.30.615695. 10.1101/2024.09.30.615695.

40. Gunn A, Stark JM. I-SceI-based assays to examine distinct repair outcomes of mammalian chromosomal double strand breaks. Methods Mol Biol Clifton NJ. 2012;920:379–91. 10.1007/978-1-61779-998-3_27.

41. Shen C, Cui T, Yang L, Gui L, Corrales-Guerrero S, Nair S, et al. KRAS-induced STN1 (OBFC1) promotes proper CTC1-STN1-TEN1 complex-independent DNA double-strand break repair and cell cycle checkpoint maintenance in pancreatic cancer. Nucleic Acids Res. 2025;53:gkaf983. 10.1093/nar/gkaf983.

42. Wickham H. ggplot2: elegant graphics for data analysis. Second edition. Cham: Springer international publishing; 2016.

43. Therneau TM, Grambsch PM. Modeling survival data: extending the Cox model. New York: Springer; 2001.

44. Liberzon A, Birger C, Thorvaldsdóttir H, Ghandi M, Mesirov JP, Tamayo P. The Molecular Signatures Database (MSigDB) hallmark gene set collection. Cell Syst. 2015;1:417–25. 10.1016/j.cels.2015.12.004.

